# Habitat preferences of specialist bat are forest-type specific: western barbastelle in natural temperate woodlands

**DOI:** 10.64898/2025.12.18.695132

**Authors:** Fabian Przepióra, Veronica Facciolati, Łukasz Janocha, Alicja Wolska, Antoni Żygadło, Michał Ciach

## Abstract

1. Understanding the species-habitat relationships of forest specialists is an essential challenge in their effective conservation. However, studies on habitat selection often neglect potential context dependence, even though species performance may differ between natural and managed stands or between forests composed of different tree species. The western barbastelle *Barbastella barbastellus*, a specialized moth predator and an old-growth forest indicator, is known to rely on fine-scale habitat structures and forest complexity, yet the relative importance of habitat traits across different forest types remains poorly understood.
2. We surveyed the species on 150 plots distributed across five the best-preserved temperate forest types in Poland: spruce, beech, beech–fir, oak–lime–hornbeam and willow–poplar forests. Bat presence was quantified using passive acoustic monitoring, and habitat characteristics were derived from LiDAR, stand inventories and surveys of tree-related microhabitats.
3. We identified canopy gaps, tree height complexity, living tree density and density of trees with insect galleries or cankers, as the best predictors of the species presence, although the strength of these relationships varied among forest types. In single-species stands, the occurrence of the western barbastelle was associated with abundance of canopy gap and complexity of tree heights, whereas in forests with more diverse tree species composition, the relationships with the stand structure were less evident and were replaced by stronger association with fine-scale habitat components such as tree-related microhabitats.
4. *Synthesis and applications.* Our study demonstrates that the effective conservation of forest specialists requires habitat-specific strategies, because no single habitat variable predicts species occurrence across contrasting forest conditions. For the western barbastelle, conservation actions should involve maintaining small canopy gaps and reduced tree height complexity in single-species stands, while preserving high heterogeneity, i.e. diverse age and size tree classes and high tree species richness in mixed forests. For a large group of forest specialist, strategies tailored to forest type, rather than a one-size-fits-all approach, should be considered.

## 1. Introduction

Habitat selection is one of the fundamental processes shaping species distributions and population dynamics (Northrup et al., 2022). Specialists, i.e. species with a narrow niche, show a particularly strong dependence on specific resources and structural attributes of ecosystems. However, increasing evidence suggests that habitat selection of a species can be context-dependent and, for example, in forest specialists may vary with forest type, stand structure, management history and environmental conditions operating across multiple spatial scales (Ettwein et al., 2020; Hendel et al., 2025; McGarigal et al., 2016; Müller et al., 2015). Understanding how forest specialists select habitat under different ecological contexts is essential for interpreting the mechanisms that drive species distributions and for designing effective conservation and management strategies across heterogeneous forest landscapes (Marini et al., 2019).

Bats provide essential ecosystem services, such as regulating insect populations and maintaining ecosystem balance (Kunz et al., 2011). Their sensitivity to alternations in tree size or age structure and tree species composition of forests makes them valuable indicators of environmental quality (Russo and Jones, 2015). The western barbastelle *Barbastella barbastellus* is one of the European bat species most sensitive to habitat alternations and is widely recognized as a forest specialist (Dietz et al., 2009). The species is distributed across much of Europe, but its range is fragmented, and thus the species is listed as “Near Threatened” on the IUCN Red List, being strictly protected under the EU Habitats Directive (Annexes II and IV; IUCN, 2016). Western barbastelle is considered specialized insectivore, with diets dominated by small- and medium-sized noctuid and geometrid moths (Carr et al., 2020; Rydell et al., 1996b; Sierro and Arlettaz, 1997). For roosting, the species selects structures occurring on dead standing trees (hereafter, snags) or mature living trees such as wood crevices, fissures and loose bark patches, while cavities – particularly woodpecker holes – are used less frequently (Rachwald et al., 2022a, 2022b; Russo et al., 2004), changing roosting sites every few days (Russo et al., 2005). Consequently, the species is strongly associated with old-growth forests that provide abundant large trees with diverse Tree-related Microhabitats (hereafter, TreMs) and high volumes of deadwood, including areas shaped by local disturbances and canopy gaps dynamic (Rachwald et al., 2022a, 2022b; Russo et al., 2004; Sierro, 1999; Toffoli and Cucco, 2020). The combination of trophic specialisation and reliance on senile forest attributes makes the western barbastelle both sensitive to alterations in forest structure and a valuable umbrella species, as its conservation safeguards an array of forest-dwelling taxa that rely on deadwood, TreMs, and heterogeneous stand conditions (Hendel et al., 2025; Russo et al., 2015). The western barbastelle is thus a suitable model organism for assessing how forest characteristics shape the occurrence of specialist species, which may in turn contribute to more effective conservation of forest biodiversity.

Previous studies have emphasized that the western barbastelle is an edge specialist, favouring semi-open conditions such as small canopy gaps, forest margins and mosaic-like landscapes (Kusch et al., 2004; Zeale et al., 2012). Yet, the importance of some structural variables has varied across studies. For instance, Froidevaux et al. (2021) demonstrated that the relative importance of tree composition is strongly context-dependent: conifers became valuable to western barbastelles in broadleaf-dominated landscapes, while broadleaves gained habitat patch importance in conifer-dominated ones. Telemetry studies showed that western barbastelle adjust their foraging activity depending on the landscape matrix: in more agricultural areas they often used riparian corridors and woodland edges, whereas in more forested landscapes they concentrated their activity within forest interiors (Carr et al., 2020a). These findings suggest that species occurrence is shaped not only by local habitat characteristics but also by landscape context and resource distribution.

Despite growing body of evidence, most research has been conducted in secondary or managed forests, where forest stand structure – i.e. tree species composition, number of age classes, vertical layering of the forest canopy, diversification of canopy closure – is simplified and saproxylic resources are impoverished (Alder et al., 2021; Froidevaux et al., 2021; Hendel et al., 2023). Consequently, little is known about how the western barbastelle respond to structural variation in forests of high naturalness, including old-growth or near-primeval stands. These systems are characterized by rich deadwood legacies, structural complexity and high invertebrate diversity (Jaroszewicz et al., 2019; Paillet et al., 2010), meaning that stand-level structural elements and food resources that are scarce or limited in managed forests may no longer constrain bats distribution in natural ones. Moreover, it remains unclear whether forest type acts as a mediator of habitat associations – that is, whether the importance of structural attributes such as canopy gaps, deadwood, or TreMs differ between single-species coniferous or deciduous stands, mixed stands and riparian forests. Addressing this gap is critical because conservation strategies based on habitat associations derived from managed landscapes may not adequately capture the requirements of western barbastelle that can be applied across all habitat contexts.

In this study, we investigated the occurrence of the western barbastelle across five forest types of high naturalness in Poland, ranging from single-species coniferous and deciduous forests to mixed and riparian systems. The species was used as a model forest specialist to explore how habitat selection varies across contrasting ecological contexts. Specifically, we tested whether forest type mediates the relationships between fine-scale habitat structure including canopy gaps, canopy height complexity, tree density, TreMs and the western barbastelle occurrence. We expected structural variables to relate differently to species occurrence across forest types, reflecting their contrasting structural and ecological complexity – particularly between simpler single-species stands and more heterogeneous, multi-species forests. By addressing these questions, our study provides the first evidence of fine-scale habitat associations of the western barbastelle in natural forests, but also explores a broader question: whether habitat selection in forest specialists is context-dependent and varies across the diversity of forest types they occupy.

## 2. Materials and methods

### 2.1. Study area

The study was conducted in five of the best-preserved natural or primeval temperate forests in Poland, characterized by minimal human impact, representing a gradient of habitats with varying levels of ecological and biological complexity, ranging from single-species coniferous or deciduous forests to mixed and multi-species forests (Fig. 1). The studied forests include Norway spruce *Picea abies* mountain forest (hereafter, spruce forest) in the Tatra Mountains (Mts.), protected as Tatra National Park (NP; established in 1954); European beech *Fagus sylvatica* mountain forest (hereafter, beech forest) in the Bieszczady Mts., protected as Bieszczady NP (established in 1973); European beech–silver fir *Abies alba* mixed mountain forest (hereafter, beech–fir forest) in Świętokrzyskie Mts., protected as Świętokrzyski NP (established in 1950); willow *Salix* sp.–poplar *Populus* sp. riparian forest (hereafter, willow–poplar forest) in the middle course of the Vistula valley, protected as Natura 2000 Special Protection Areas; and pedunculate oak *Quercus robur*–small-leaved lime *Tilia cordata*–European hornbeam *Carpinus betulus* forest (hereafter, oak–lime–hornbeam forest) in Białowieża forest, protected as Białowieża NP (established in 1932). All selected forests are either under strict conservation regime or, in the case of riparian forest, located within the river channel, which is hardly accessible. Hence, in all selected habitats forestry operations such as tree removal, planting and logging are either prohibited or absent, and the stands have developed and persisted through natural processes without human intervention. Due to their pristine nature, the studied forests in Bieszczady NP and Białowieża NP are protected as UNESCO World Heritage sites (Kujawa et al., 2016; UNESCO, 2021).

**Fig. 1.**
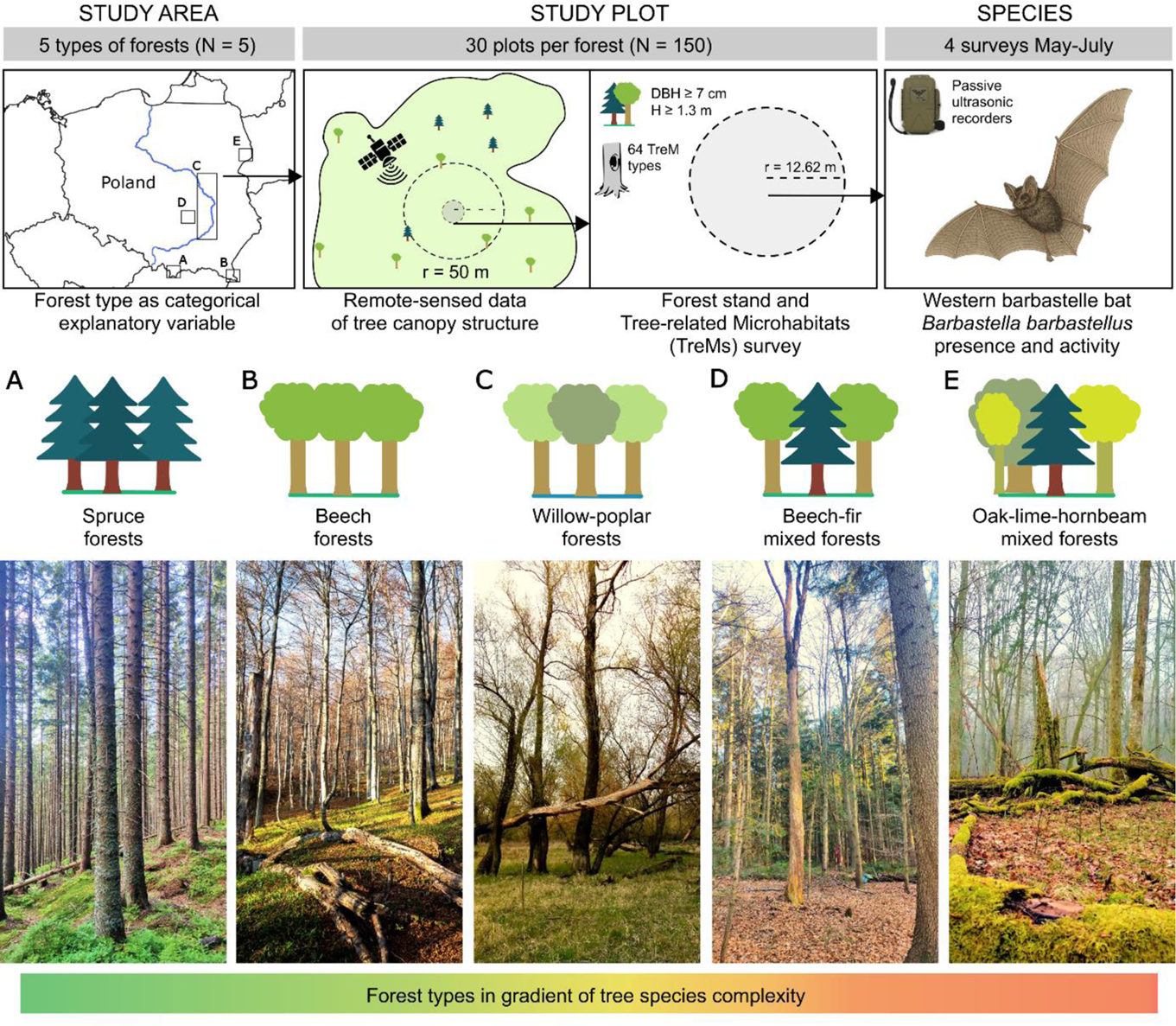
Study area, sampling design and investigated forest types. Studied forests in Poland included Tatra National Park (NP) (A), Bieszczady NP (B), Middle Vistula Valley (C), Świętokrzyski NP (D) and Białowieża NP (E). Within each forest, 30 study plots (N = 150 in total) were established, where tree canopy structure was quantified using airborne LiDAR and field surveys of stand attributes and Tree-related Microhabitats (TreMs; DBH – diameter at breast height (∼1.3 m); H – height of a tree; “TreM types” refers to the structures listed in Kraus et al. (2016)). The western barbastelle *Barbastella barbastellus* presence and activity were monitored using passive ultrasonic detectors during four full-night surveys per plot in May–July. Panels (A–E) illustrate the five studied forest types: Norway spruce *Picea abies* forest, European beech *Fagus sylvatica* forest, willow *Salix* spp.–poplar *Populus* spp. Riparian forest, European beech–silver fir *Abies alba* mixed forest, and pedunculate oak *Quercus robur*–small-leaved lime *Tilia cordata*–European hornbeam *Carpinus betulus* mixed forest (photos by Fabian Przepióra).

The studied forests exhibit an average number of tree species ranging from 1.1 ±0.4 SD species per plot in beech forest (Przepióra et al., 2025), up to 4.0 ±1.0 SD species per plot in oak–lime–hornbeam forest (Przepióra and Ciach, 2023). In the coniferous forest in Tatra Mts., Norway spruce dominates with admixture of rowan *Sorbus aucuparia* (Bodziarczyk et al., 2019), while in the mountainous deciduous forest in Bieszczady Mts., European beech is dominant (Michalik and Szary, 2016). In Świętokrzyski NP, silver fir dominates with admixtures of European beech and rowan and Norway spruce (Orzechowski and Kędziora, 2023). In Białowieża Forest, European hornbeam, small-leaved lime, Norway spruce and pedunculate oak dominate (Keczyński, 2017). The willow–poplar forest in the Middle Vistula Valley is dominated by crack willow *Salix fragilis*, white willow *Salix alba*, black poplar *Populus nigra* and white poplar *Populus alba* (Przepióra and Ciach, 2022).

The studied forests are characterized by ecological continuity, i.e. forest communities have persisted without major interruption since the last glaciation (Nordén and Appelqvist, 2001), with deadwood resources represented by various forms, such as snags, fallen logs and stumps, all in different stages of decay, ranging from 45.8 m³ ha^-1^ in beech–fir forest (Figarski et al., 2014), 55.0 m³ ha^-1^ in beech forest (Kacprzyk et al., 2014), 159.0 m³ ha^-1^ in oak–lime– hornbeam forest (Bobiec, 2002), to 248.0 m³ ha^-1^ in spruce forest (Bodziarczyk et al., 2019). These forests also feature exceptionally rich TreM resources, with between 49 and 61 different TreM types found across the studied areas (Przepióra and Ciach, 2025, 2023). Due to the strict conservation regime, natural processes such as wind disturbances, cambium-feeding insect activity, river floods and lightning strikes occur spontaneously, increasing habitat heterogeneity (Jaroszewicz et al., 2019; Przepióra and Ciach, 2022).

Bat assemblages in the studied forests have species richness ranging from 10 to 19 species. In montane beech or spruce stands, assemblages are dominated by small *Myotis* spp. bats (Piksa et al., 2022; Sachanowicz and Wower, 2013), while diversity peaks in mixed and lowland broadleaf forests, where both open-space foragers (e.g. *Nyctalus noctula*, *Pipistrellus* spp.) and forest specialists are present (Dietz et al., 2018; Rachwald et al., 2021). Across sites, the western barbastelle was recorded consistently but with varying frequency: it was the least common in high-elevation spruce or beech forests (Piksa et al., 2022; Sachanowicz and Wower, 2013), while locally common in beech–fir and oak–lime–hornbeam stands (Ciechanowski et al., 2023; Rachwald et al., 2021). The species forms a constant, though never dominant, component of Central European forest bat assemblages, with its abundance strongly influenced by study region and habitat type (Sachanowicz et al., 2006).

### 2.2. Field methods and remote sensed-data

In each of the studied forests, 30 circular plots with a radius of 12.62 m (0.05 ha; N = 150 in total) were randomly selected (Fig. 1) to maintain a minimum distance between the plots greater than 150 m. The mean distance between the ultimately selected adjoining plots in a given forest was 221.9 m ± 44.7 SD (range 161.8–406.7 m). Habitat surveys were conducted once per plot during the leaf-off seasons: from October to May in deciduous or mixed forests, and from August to September in spruce forest, during the years 2019–2024. On each study plot, all living trees and snags with DBH ≥ 7 cm were recorded (Fig. 1). For each tree, species identity, DBH, azimuth and distance from plot centre and live status (living tree or snag) were noted. Each tree was also examined for the presence of TreMs, following the standardized classification system for temperate forests (Kraus et al., 2016). To ensure consistency, all TreM inventories were conducted by a single observer, with a minimum of 3 min spent examining each tree, including the crown and upper trunk using binoculars. Every piece of coarse woody debris (CWD) ≥ 100 cm in length and ≥ 7 cm in diameter at the thinner end was measured.

Canopy structure metrics were derived from LiDAR data within a 50-m radius buffer centred on each study plot (Fig. 1). Canopy gaps were automatically identified in the Canopy Height Model (0.5-m resolution) (Silva et al., 2019), with size thresholds of 0.01–5.0 ha and a minimum canopy height difference of 3 m, ensuring detection of both small and large gaps characteristic of the studied forests (Lewandowski et al., 2021). The number of gaps intersecting the 50-m buffer was then calculated for each plot. The entropy of height distribution was computed as Shannon entropy of LiDAR point returns across vertical bins, representing canopy height complexity, with low values indicating single-layered and high values indicating multi-layered stands (Carrasco et al., 2019; Roussel et al., 2020).

Bat presence and activity was surveyed at each of the 150 study plots using Titley Chorus passive ultrasonic detectors (Titley Scientific, Brendale, QLD, Australia; Fig. 1). The survey was conducted between May and July 2022, with four recording sessions per plot carried out. During each session, detectors were deployed for a minimum of one night. If night was affected by rainfall or strong wind, the devices were left in place for an additional night to ensure data quality. During subsequent visits, detectors were mounted on the same tree at a height of up to 2 m, on the thinnest suitable tree located as close as possible to the centre of each plot. The mean DBH of the trees on which the devices were mounted was 19.3 cm ± 7.8 SD (range 7.0–39.0 m), and they were located at a mean distance of 3.9 m ± 2.1 SD (range: 0.3–11.1 m) from the centre of the study plot. Recordings of 10-min duration were made in continuous mode, starting 30 min before sunset and ending 30 min after sunrise, thus capturing full-night acoustic activity within the 10–140 kHz frequency range at a sampling rate of 500 ksps.

### 2.3. Acoustic data processing

All recordings were processed using Kaleidoscope Pro software (version 5.1.9h Wildlife Acoustics, Inc., Maynard, MA, USA). 10-min recordings (N = 33 759) were first segmented into 5-s clips (N = 4 295 642), after which those containing high levels of background noise or interference were automatically filtered out (N = 4 191 347; 97.6% of all 5-s clips). The remaining “no-noise” clips (N = 104 295; 2.6%) were then analyzed using the Auto-ID function to detect the presence of 22 bat species known to occur in the study areas (Okarma, 2024). The Auto-ID algorithm works by scanning each 5-s clip for ultrasonic signals characteristic of the selected species, assigning a species identity along with a confidence score. Clips containing signals that could not be confidently attributed to any known bat species were labelled “no-ID”, while clips identified as containing non-bat sounds (e.g., insects, wind, mechanical noise) were labelled “noise”. After this stage, the dataset included 54 941 clips with acoustic detections attributed to 22 bat species/taxa, 23 874 no-ID clips and 25 480 noise clips. To validate the accuracy of the automated classification, 842 clips assigned to western barbastelle, 795 clips assigned to other bat species, and 1000 noise and no-ID clips were manually reviewed. Manual validation confirmed that 694 (82.4%) of all clips initially assigned to the western barbastelle were correctly identified, and 23 (2.9%) clips initially assigned to other bat species were in fact western barbastelle. Among the noise clips, 728 contained only non-bat sounds, while the remaining included bat calls that had been misclassified. For no-ID clips, 772 contained genuine bat calls, although these could not be reliably assigned to any species.

### 2.4. Data handling and analyses

The species prevalence, i.e. the number of study plots with species occurrence recorded at least on one visit (N), the species activity, i.e. the highest number of 5-s long clips with recorded species ultrasound pulses recorded within one visit (5-s clips per night) and recording effort expressed as total number of 10-min long continuous ultrasonic recordings obtained with all four visits (10-min recordings), on each study plot were calculated. Habitat characteristics describing the canopy structure, forest stand structure and composition, and TreM assemblage was calculated for each plot, and the mean, standard error (SE) and range of these characteristics were calculated based on all 150 study plots. The canopy structure included number of canopy gaps (N) and canopy height complexity (value 0–1; single- to multi-layered stands). Forest stand structure characteristics included the density (trees ha^-1^) and mean DBH (cm) of measured trees calculated separately for living trees and snags, and the diversity of tree species recorded on the plot (Shannon-Wiener index) and total volume of CWD found on the plot (m^3^). The characteristics of the TreM assemblage calculated on the basis of all inventoried TreM types included two TreM indices: TreM density, i.e. the total density of trees bearing TreMs (if a tree bore several TreM types, it was counted once for each TreM type) extrapolated to one ha (expressed as TreM-bearing trees ha^-1^) and TreM diversity, i.e. the Shannon–Wiener diversity index calculated on the basis of TreM types recorded and their relative abundance.

For each plot, the density of TreM-bearing trees was calculated separately for each recorded TreM type. This measure represents the number of trees bearing a given TreM type extrapolated to 1 ha. Due to potential relevance of TreMs to the occurrence and activity of the western barbastelle, selected TreM types were assigned to eleven TreM groups by summing their densities: (i) woodpecker cavities, (ii) trunk and mould cavities with ground contact, (iii) trunk and mould cavities without ground contact, (iv) large semi-open trunk and mould cavities, (v) branch holes, (vi) insect galleries and bore holes, (vii) cracks and crevices, (viii) bark pockets, (ix) cankers and burrs, (x) canopy deadwood and (xi) fruiting bodies of fungi (Table S1).

Differences in mean of canopy structure characteristics, tree structure, TreM assemblages and the densities of TreM groups between study plots located in different forest types were tested using ANOVA, followed by Tukey’s post hoc test. The relationships between habitat characteristics and species occurrence were analysed using generalized linear models (GLMs) with a binomial error distribution and a logit link function. The modelling process began with a null model including only the intercept. This was followed by a model containing a single categorical variable representing forest type, defined by five levels: spruce, beech, beech–fir, willow–poplar, and oak–lime–hornbeam. Subsequently, a series of models were constructed that combined forest type with time effort and different sets of continuous explanatory variables, each corresponding to a distinct group of ecological or structural characteristics. These comprised canopy-related variables, including the number of canopy gaps and canopy height complexity, tree structure parameters such as tree species diversity, the density of living trees, density of snags, DBH of both living trees and snags, volume of CWD, TreM indices, i.e. TreM density and TreM diversity, and TreM group-level variables including all eleven TreM groups (Table S1) For each model, backward selection was applied by sequentially removing non-significant variables from the initial full model, with the final model determined based on the lowest Akaike Information Criterion (AIC). In total, six models were obtained: an intercept-only model, a model with forest type alone and four models including forest type along with one of the respective groups of continuous variables. None of the final models showed evidence of residual overdispersion or inflation (p = 0.832–0.920 and p = 1.0–1.0, respectively).

To test for differences in model estimates between forest types, we performed post-hoc pairwise comparisons of estimated marginal means. Pairwise contrasts of estimates were extracted for all combinations of forest types. These contrasts quantify whether the predicted probability of western barbastelle occurrence differs significantly between forest types, after accounting for habitat covariates included in the models. All statistical procedures were performed using R 3.5.0 software (CoreTeam R., 2017). The GLM analyses and post hoc tests were carried out using the glmmTMB and emmeans packages, while LiDAR-derived metrics and canopy gaps detection were computed with the lidR and ForestGapR packages, respectively (Lenth, 2025; Magnusson et al., 2017; Roussel et al., 2020; Silva et al., 2019).

## 3. Results

The western barbastelle was recorded on 45% of all study plots (N = 150), with the highest prevalence observed in mixed forest – 77% of plots (N = 30) in oak–lime–hornbeam forest and 60% (N = 30) in beech–fir forest – followed by 40% (N = 30) in spruce forest, 30% (N = 30) in willow–poplar forest and 17% (N = 30) in beech forest. Mean activity of the species across all survey nights was 3.6 ± 0.9 5-s clips per plot and did not differ between forest types (Fig. 2). Recording effort did not vary between forest types, with an overall mean of 239.3 ± 3.3 10-min recordings per plot (Fig. 2).

**Fig. 2.**
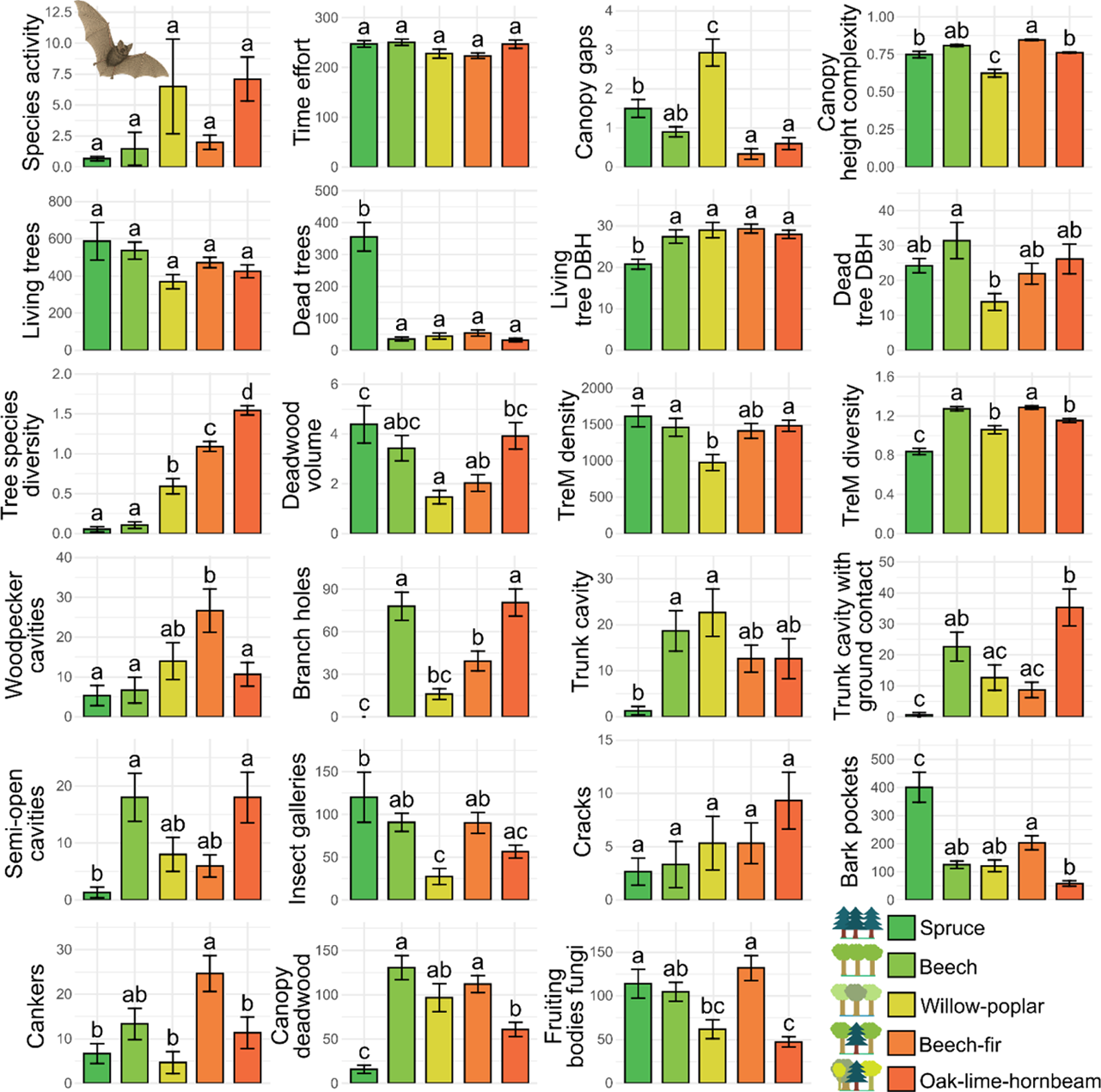
Mean (± SE) activity of the western barbastelle *Barbastella barbastellus* (the highest number of 5-s clips containing species ultrasound pulses recorded within a single visit) and sampling effort (total number of 10-min recordings per plot), along with habitat characteristics measured at the study plots. Variables included: canopy-related (number of canopy gaps; canopy height complexity), stand structure-related (density of living and dead standing trees, trees ha⁻¹; diameter at breast height (DBH) of living and dead trees, cm; tree species diversity, Shannon–Wiener index; volume of lying deadwood, m³), and variables describing Tree-related Microhabitats (TreMs) (TreM density, TreM-bearing trees ha⁻¹; TreM diversity, Shannon–Wiener index; and the density of specific TreM groups: woodpecker cavities; branch holes, trunk and mould cavities with and without ground contact; semi-open cavities; insect galleries and boreholes; cracks; bark pockets; cankers; canopy deadwood; and fungal fruiting bodies, all expressed as TreM-bearing trees ha⁻¹). Results are shown for plots located in Norway spruce *Picea abies* forest (N = 30), European beech *Fagus sylvatica* forest (N = 30), European beech–silver fir *Abies alba* mixed forest (N = 30), pedunculate oak *Quercus robur*–small-leaved lime *Tilia cordata*–European hornbeam *Carpinus betulus* mixed forest (N = 30) and willow *Salix* spp.–poplar *Populus* spp. riparian forest (N = 30). Differences in mean values between forest types were tested using ANOVA with Tukey’s post hoc test; letter indices (a–c) indicate significance at p < 0.05.

Tree species diversity followed the expected gradient across forest types, being highest in oak–lime–hornbeam forest, followed by beech–fir and riparian forest, and lowest in beech and spruce stands (Fig. 2). The forest types differed in the number of canopy gaps, which was highest in riparian forest and lowest in oak–lime–hornbeam and beech–fir stands (Fig. 2). Canopy height complexity was the lowest in riparian forest and the highest in beech–fir and beech forests (Fig. 2). The density and mean DBH of living trees were similar among forest types, except in spruce forest, which supported markedly higher densities of snags and smaller DBH of living trees compared with the other forest types (Fig. 2). Snag DBH, the volume of CWD and TreM density differed between the most contrasting habitats (Fig. 2). TreM diversity grouped into clusters: beech and beech–fir forests had the highest and most similar values, oak–lime–hornbeam and riparian forests held intermediate values, and spruce forest showed the lowest TreM diversity (Fig. 2). The density of individual TreM types varied among forest types, reflecting habitat-specific TreM assemblages (Fig. 2). For instance, spruce forest contained numerous bark pockets but few branch holes or trunk cavities, whereas the latter reached their highest values in beech and oak–lime–hornbeam forest (Fig. 2).

The model combining forest type with canopy-structure variables provided the best explanation for the presence of the western barbastelle (Table 1). The second-best model included forest type and forest-structure variables, followed by the model including forest type and TreM groups (Table 1). Several habitat characteristics significantly influenced the presence of the western barbastelle: for canopy-related variables, the probability of species occurrence increased with the number of canopy gaps and decreased with canopy height complexity (Table 2a). For tree structure, occurrence was negatively related to the density of living trees, while the density of snags showed a weak, near-significant positive association (Table 2b). For TreM indices, TreM diversity showed a marginal but positive association with the presence of the western barbastelle (Table 2c). For specific TreM groups, occurrence increased with the density of insect galleries and cankers, while the association with trunk and mould cavities with ground contact was marginally negative (Table 2d).

**Table 1.** Model selection results for generalised linear models with a binomial distribution describing the occurrence of the western barbastelle *Barbastella barbastellus*. Candidate models included a null model (intercept only), forest type alone (a categorical variable with five levels), and forest type combined with different sets of continuous predictors (canopy structure, tree structure, Tree-related Microhabitat (TreM) indices, and TreM groups). Models were ranked according to Akaike’s Information Criterion (AIC), with lower AIC values indicating better fit; ΔAIC represents the difference relative to the best model.

| Model of species occurrence | AIC | ΔAIC |
| --- | --- | --- |
| Null model | 208.2 | 47.2 |
| Forest type | 187.0 | 26.0 |
| Forest type + Canopy structure | 161.0 | 0 |
| Forest type + Tree structure | 172.4 | 11.4 |
| Forest type + TreM indices | 185.6 | 24.6 |
| Forest type + TreM groups | 177.3 | 16.3 |

**Table 2.** Final models describing the relationship between the occurrence of the western barbastelle *Barbastella barbastellus* and habitat characteristics included as numeric predictors: **(a)** canopy-related factors – number of canopy gaps and canopy height complexity; **(b)** tree structure-related factors – density of living trees and density of dead standing trees; **(c)** Tree-related Microhabitat (TreM) indices – TreM density and TreM diversity; and **(d)** specific TreM groups – density of branch holes, trunk and mould cavities with ground contact; insect galleries and boreholes; bark pockets; and cankers. Models were fitted for study plots located in Norway spruce *Picea abies* forest (N = 30), European beech *Fagus sylvatica* forest (N = 30), European beech–silver fir *Abies alba* mixed forest (N = 30), pedunculate oak *Quercus robur*–small-leaved lime *Tilia cordata*–European hornbeam *Carpinus betulus* mixed forest (N = 30) and willow *Salix* spp.–poplar *Populus* spp. riparian forest (N = 30). Forest type was included as a categorical factor in all models, with oak–lime– hornbeam mixed forest as the reference level.

| Models of species occurrence | Estimate | SE | Z | P |
| --- | --- | --- | --- | --- |
| **(a) Canopy structure** |  |  |  |  |
| Intercept | 12.13 | 3.35 | 3.62 | 0.000 |
| Number of canopy gaps | 0.40 | 0.19 | 2.14 | 0.033 |
| Canopy height complexity | -14.60 | 4.34 | -3.36 | 0.001 |
| Beech forest | -2.42 | 0.69 | -3.51 | 0.000 |
| Willow–poplar forest | -6.08 | 1.38 | -4.40 | 0.000 |
| Beech–fir forest | 0.55 | 0.69 | 0.80 | 0.426 |
| Spruce forest | -2.20 | 0.69 | -3.18 | 0.001 |
| **(b) Tree structure** |  |  |  |  |
| Intercept | 2.23 | 0.58 | 3.84 | 0.000 |
| Density of living trees | 0.00 | 0.00 | -3.20 | 0.001 |
| Density of dead trees | 0.00 | 0.00 | 1.86 | 0.063 |
| Beech forest | -2.69 | 0.67 | -4.04 | 0.000 |
| Willow–poplar forest | -2.33 | 0.61 | -3.83 | 0.000 |
| Beech–fir forest | -0.80 | 0.58 | -1.37 | 0.170 |
| Spruce forest | -2.87 | 1.00 | -2.88 | 0.004 |
| **(c) TreM indices** |  |  |  |  |
| Intercept | -1.53 | 1.58 | -0.97 | 0.334 |
| TreM diversity | 2.37 | 1.34 | 1.78 | 0.076 |
| Beech forest | -3.14 | 0.69 | -4.53 | 0.000 |
| Willow–poplar forest | -1.90 | 0.60 | -3.18 | 0.001 |
| Beech–fir forest | -1.12 | 0.61 | -1.84 | 0.066 |
| Spruce forest | -0.88 | 0.69 | -1.27 | 0.206 |
| **(d) TreM groups** |  |  |  |  |
| Intercept | 0.66 | 0.70 | 0.95 | 0.342 |
| Branch holes | 0.01 | 0.01 | 1.53 | 0.127 |
| Trunk cavity with ground contact | -0.02 | 0.01 | -1.73 | 0.083 |
| Insect galleries | 0.01 | 0.00 | 2.47 | 0.014 |
| Cankers | 0.03 | 0.01 | 2.10 | 0.036 |
| Beech forest | -3.72 | 0.79 | -4.69 | 0.000 |
| Willow–poplar forest | -1.75 | 0.74 | -2.37 | 0.018 |
| Beech–fir forest | -1.53 | 0.76 | -2.02 | 0.043 |
| Spruce forest | -1.94 | 0.82 | -2.36 | 0.018 |

Predicted response curves (Fig. 3) and pairwise contrasts (Fig. 4) showed that the strength of relationships between habitat characteristics and species occurrence differed across forest types. The relationship between the occurrence of the western barbastelle and canopy structure variables was evident in single-species forests (spruce or beech), but weak in mixed forests (beech–fir or oak–lime–hornbeam) and in riparian forest (willow–poplar) (Fig. 3). Within single-species forests and within mixed forests, predicted species occurrence did not differ, whereas the probability of species occurrence between these groups differs at comparable levels of canopy structure variables, and riparian forest differed from both these forest types (Fig. 4). The relationship between species occurrence and the density of living tree was most evident in mixed forests, whereas the association with snag density was most evident in single-species stands (Fig. 3). At comparable levels of tree structure variables, predicted species occurrence differed significantly between both mixed forests and each of the single-species forests, as well as between both mixed forests and riparian forest (Fig. 4). The relationships between species occurrence and TreM diversity were most evident in spruce, mixed and riparian forests (Fig. 3), and the largest differences in predicted occurrence at comparable levels of TreM diversity were between beech and mixed forests, and between mixed and riparian forests (Fig. 4). The relationship between species occurrence and the density of trunk and mould cavities with ground contact was most evident in oak–lime– hornbeam forest (Fig. 3), whereas the relationship between species occurrence with the density of insect galleries and boreholes, or the density of cankers, were most evident in spruce, beech–fir and riparian forests (Fig. 3). The largest differences in predicted probabilities of species occurrence at comparable TreM densities occurred between single-species and mixed stands, and between single-species and riparian forests (Fig. 4).

**Fig. 3.**
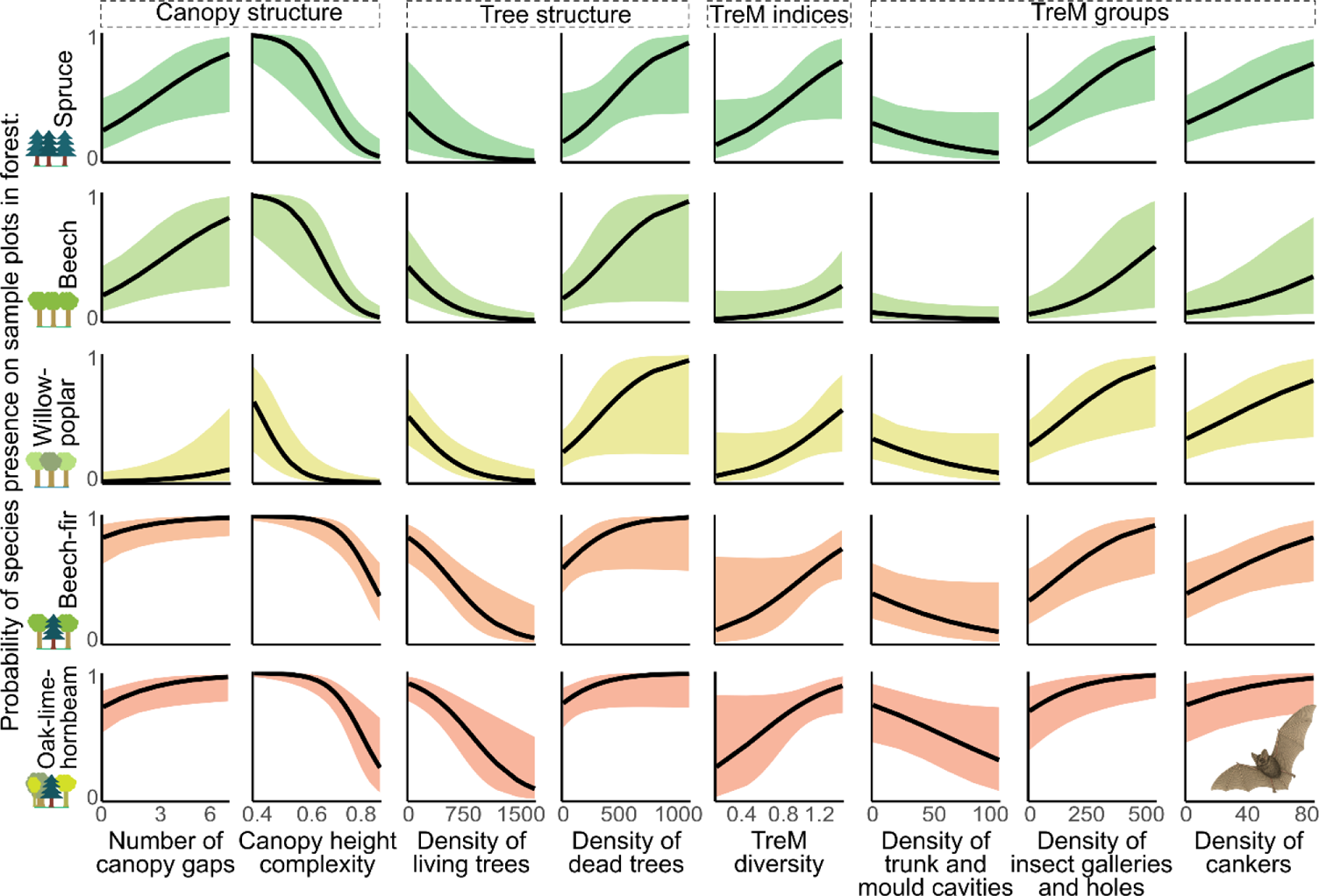
Relationships between the probability of occurrence of the western barbastelle *Barbastella barbastellus* and habitat characteristics identified as significant or near-significant in the final models (see Table 2). Variables included: canopy-related (number of canopy gaps; canopy height complexity), tree structure-related (density of living and dead standing trees, trees ha⁻¹), Tree-related Microhabitat (TreM) indices (TreM diversity, Shannon–Wiener index), and TreM groups (density of trunk and mould cavities with ground contact; insect galleries and boreholes; bark pockets; and cankers, all expressed as TreM-bearing trees ha⁻¹). Results are shown for study plots located in Norway spruce *Picea abies* forest (N = 30), European beech *Fagus sylvatica* forest (N = 30), European beech–silver fir *Abies alba* mixed forest (N = 30), pedunculate oak *Quercus robur*–small-leaved lime *Tilia cordata*–European hornbeam *Carpinus betulus* mixed forest (N = 30) and willow *Salix* spp.–poplar *Populus* spp. riparian forest (N = 30). Coloured ribbons indicate 95% confidence intervals.

**Fig. 4.**
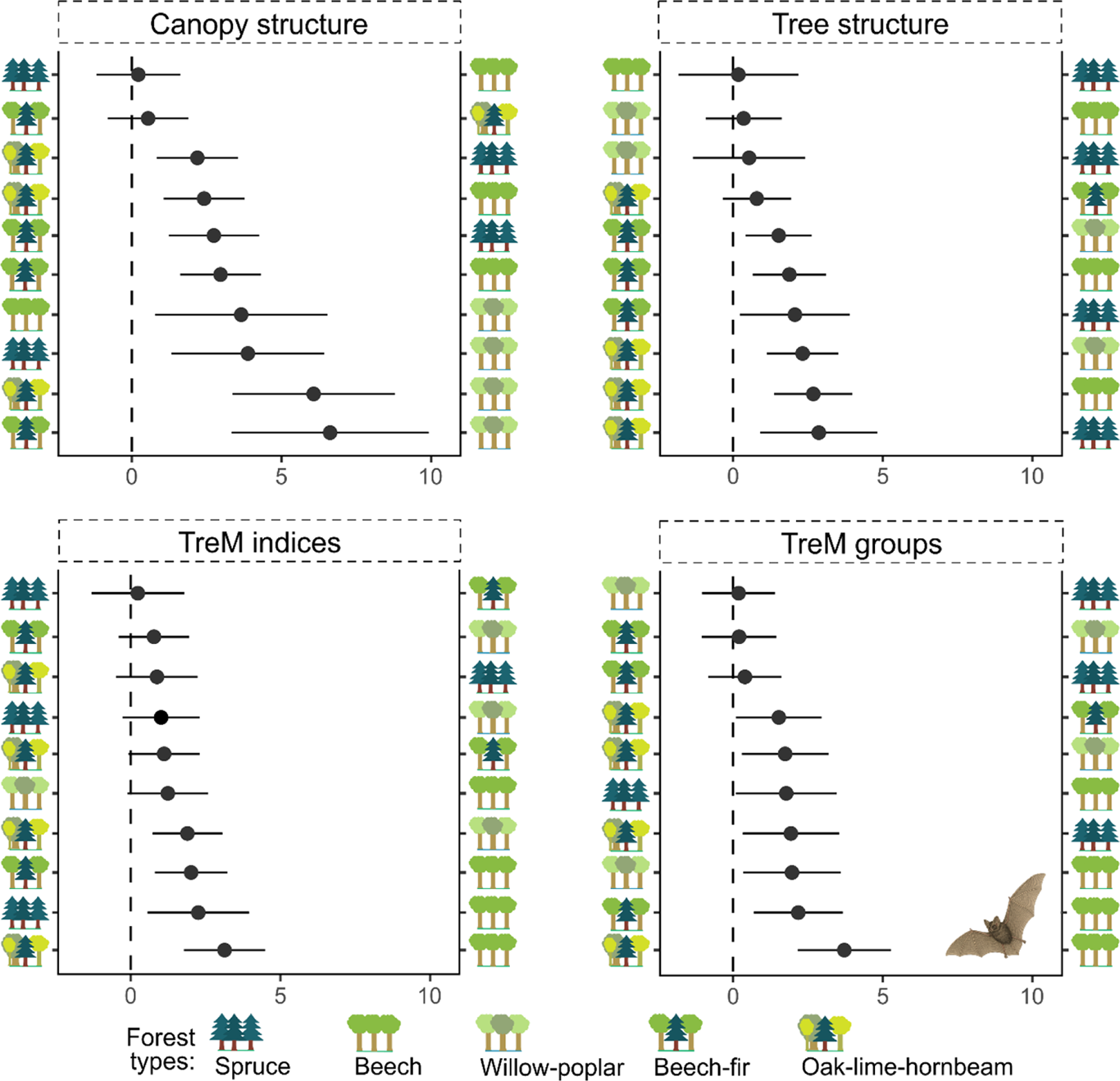
Pairwise contrasts of the estimated probability of occurrence of the western barbastelle *Barbastella barbastellus* between forest types (logit scale; left–right estimates subtracted): Norway spruce *Picea abies* forest (N = 30), European beech *Fagus sylvatica* forest (N = 30), European beech–silver fir *Abies alba* mixed forest (N = 30), pedunculate oak *Quercus robur*– small-leaved lime *Tilia cordata*–European hornbeam *Carpinus betulus* mixed forest (N = 30) and willow *Salix* spp.–poplar *Populus* spp. riparian forest (N = 30). Models were based on averaged habitat predictors, including canopy and tree structure, and Tree-related Microhabitat (TreM) indices and TreM groups (see Table 2 and Fig. 3). Confidence intervals (solid lines) that include zero denote non-significant differences.

## 4. Discussion

Our modelling indicated that canopy and tree structure were the main drivers of the occurrence of the western barbastelle, but the strength of these relationships differed among forest types. The response curves grouped the studied forests into three functional categories: (i) single-species (spruce or beech), (ii) mixed deciduous–coniferous (beech–fir and oak– lime–hornbeam), and (iii) riparian willow–poplar forest. In single-species forests, relationships such as a positive association with canopy gap abundance or a negative association with high canopy height complexity were most pronounced, while in mixed or riparian forests, relationships between the probability of occurrence of the western barbastelle and canopy structure were less evident. The differences in habitat association among forest types can be explained by species ecology and resource availability across the studied systems (Erasmy et al., 2021; Hendel et al., 2023; Russo et al., 2025, 2004; Toffoli and Cucco, 2020). The western barbastelle is a dietary specialist, feeding almost exclusively on nocturnal moths (Rydell et al., 1996b; Sierro and Arlettaz, 1997), and its low-amplitude echolocation calls – adapted to detect tympanate moths (Seibert et al., 2015) – reduce detection range, which constrains foraging efficiency in open habitats (Zeale et al., 2012). Dense stands with closed canopy are suboptimal as they provide low abundances of small- and medium-sized moths and limit manoeuvrability (Froidevaux et al., 2021; La Cava et al., 2024), whereas small gaps and a single canopy layer enhance prey activity and warm subcanopy temperatures (Forrester et al., 2012), lowering thermoregulatory demands for bats emerging from roosts (Russo et al., 2007; Rydell et al., 1996a). Thus, even within natural forests, the western barbastelle selects patches with locally reduced vertical and horizontal canopy closure, and reliance on such canopy openings is strongest in single-species forests, which are typically more closed and provide fewer suitable foraging opportunities than mixed or riparian forests with high habitat heterogeneity.

Forest type dependence of the strength and the direction of relationships between forest structural parameters and occurrence of forest specialists align with the habitat complementation hypothesis (Dunning et al., 1992), which posits that the relative importance of habitat components depends on the availability of complementary resources across the landscape. In mixed forests, the western barbastelle can exploit a broad range of habitat patches and foraging strategies, reducing the relative importance of individual structural elements. In riparian forest, likewise, high environmental dynamics, access to water and elevated abundances of prey associated with seasonal pulses, e.g. flowering willows in early spring or the mass emergence of water-related species, can override the importance of individual structural elements (Apoznański et al., 2023; Carr et al., 2023; Zeale et al., 2012), leading to weaker associations with small-scale habitat variables (Hendel et al., 2023). Hence, structural habitat elements in simple stands lose their importance in complex multi-species forests, where greater habitat complexity provides alternative resources. However, the forest-type-specific habitat associations in the western barbastelle need to be investigated in other specialists with narrow trophic or roosting requirements.

Our results indicate that snag density was only a weak predictor of the occurrence of the western barbastelle occurrence in forests of high naturalness. A likely explanation is that deadwood resources were exceptionally abundant and structurally diverse across all study plots, creating an insufficient gradient to detect clear effects at fine scales. In contrast to managed forests, where snags often constitute a limiting resource for many bat species, in natural stands this resource may be so ubiquitous that its predictive power becomes diluted (Alder et al., 2021; Tillon et al., 2016). Higher occurrence of the western barbastelle has been reported in natural forest patches with locally elevated snag densities, for example after bark beetle outbreaks in mixed stands with spruce (Rachwald et al., 2022a, 2022b). A similar pattern emerged in our study: spruce forest containing habitat patches shaped by bark beetle activity was the only forest type that showed distinct snag accumulation and exhibited an evident (though non-significant) relationship between the probability of occurrence of the species and snag density. In this context, our results show that, unlike in managed forests where deadwood scarcity makes snags key predictors of the occurrence of the western barbastelle, in natural forests this relationship weakens and remains evident only in habitats with the most pronounced gradients in deadwood abundance.

In our study, overall TreM diversity was a weak predictor of the occurrence of the western barbastelle in natural forests. This aligns with the species’ ecology: a specialised predator that uses only a few TreM types as roosts and forages on prey associated with specific TreMs. Accordingly, the most pronounced associations emerged with saproxylic TreMs such as insect boreholes and cankers, with the relationship between their densities and the probability of occurrence of the western barbastelle being most evident in forests with lower tree species diversity. While most identified prey moths are phytophagous (Apoznański et al., 2023; Carr et al., 2020; Rydell et al., 1996b), these TreMs host rare moth species whose larvae develop in decaying wood or fungi, as well as other invertebrates that may supplement the species’ diet (Jaworski et al., 2016, 2014; Larrieu et al., 2018). The western barbastelle is known to exploit local and seasonal prey peaks, for example by feeding on Orthosia moths attracted to flowering willows in early spring (Apoznański et al., 2023). In natural forests, foraging flexibility may favour patches bearing TreMs that support not only common phytophagous taxa but also saproxylic species (Bury et al., 2014), with such patterns being more pronounced in forests with low structural complexity. While previous studies in secondary or managed forests, where saproxylic fauna is relatively impoverished (Müller and Bütler, 2010), reported only weak or moderate bat–TreM associations (Basile et al., 2020; Paillet et al., 2018), our results from natural stands reveal a stronger link between species occurrence and the abundance of specific TreM types. This indicates that under natural forest conditions, some saproxylic TreMs may represent critical microhabitats in certain forest types, supporting additional prey resources for the western barbastelle.

In natural forests, trees bearing trunk and mould cavities with ground contact were weak negative predictors of the occurrence of the western barbastelle, with the relationship being most pronounced in highly diverse stands such as the Białowieża Forest. The western barbastelle prefers warm, dry and elevated roosts that reduce humidity fluctuations and predation risk (Dietz et al., 2018; Russo et al., 2015, 2004), whereas basal cavities are typically cooler, more humid and more accessible to ground-dwelling predators (Larrieu et al., 2018). In the Białowieża Forest, where the densities of predators such as pine marten *Martes martes* are high (Zalewski et al., 1995), such ground-level cavities may either serve as shelters for these predator or attract them to such habitat patches during foraging. Hence, the observed negative relationship between the occurrence of the western barbastelle and density of trees bearing mould cavities with ground contact may indicate an anti-predator response, in which the western barbastelle avoids sites rich in this TreM type and prefers safer, elevated roosting structures that provide reduced exposure to predation.

Our findings are consistent with previous work demonstrating that the habitat associations of the western barbastelle are context dependent. For instance, different tree species have been identified as key predictors in different regions, with beech being particularly important in the Alps, whereas oak was more relevant in the Apennines (Russo et al., 2025). Similarly, the role of large trees varied with forest management: their presence was important in coppiced systems, where such structures were scarce, whereas in more structurally diverse stands, the number of large trees was not significant (Alder et al., 2021). Moreover, Froidevaux et al. (2021) showed that coniferous trees are valuable within deciduous-dominated landscapes, while deciduous trees are valuable within coniferous-dominated ones. Taken together, these results underline that the ecological drivers of the occurrence of the western barbastelle are not universal but that habitat–occurrence relationships depend on both stand composition and landscape context. Our study shows that even under natural conditions, forest type continues to shape the strength of species–habitat associations. As similar context-dependent patterns are expected for other forest specialists reliant on old-growth attributes, forest type should be incorporated into conservation planning for such species.

## 5. Conclusion

Our study shows that the occurrence of the western barbastelle in natural forests is driven by the interaction between forest structure and forest type. In compositionally simple stands, the species is strongly constrained by structural conditions such as canopy openness and complexity, as well as tree density, whereas in forest with higher tree species diversity these constraints are less evident and other ecological drivers, such as seasonal resource pulses or predation risk, may gain importance. Associations between the occurrence of the western barbastelle with deadwood or TreMs were generally limited, yet saproxylic features such as insect galleries and cankers emerged as relevant habitat predictors in certain forest types. Our findings demonstrate that habitat selection in forest specialists is context dependent and therefore cannot be generalised across forest types. From a conservation perspective, our results suggest that the effective protection of forest specialist species requires strategies tailored to forest type rather than a one-size-fits-all approach. Using the western barbastelle as a model species, we highlight that maintaining small canopy gaps and low canopy height complexity is crucial for supporting species presence in single-species forests, while preserving heterogeneous tree size structure and diverse tree species composition is key in mixed stands. This principle is likely to apply across forest specialists that depend on features typical of old-growth forests.

## Authors’ contributions

Michał Ciach and Fabian Przepióra conceived the idea and designed the methodology. Fabian Przepióra, Łukasz Janocha, Veronica Facciolati, Alicja Wolska and Antoni Żygadło collected and annotated the acoustic data. Fabian Przepióra analysed the data. Michał Ciach and Fabian Przepióra led the writing of manuscript. All authors contributed critically to the draft and gave final approval for publication.

## Acknowledgements

This study was financially supported by The National Science Centre in Poland from a Preludium grant (2021/41/N/NZ9/03441) and an Opus grant (2021/41/B/NZ8/03456), and also by the Ministry of Science and Higher Education of the Republic of Poland within the framework of statutory funds awarded to the Faculty of Forestry, University of Agriculture. Fabian Przepióra was supported by the Foundation for Polish Science (FNP). We thank the management of the Tatra National Park, Bieszczady National Park, Świętokrzyski National Park and Białowieża National Park for permission to carry out. We would also like to thank Arkadiusz Fröhlich, Damian Kurlej, Izabela Fedyń and Jakub Wyka for their assistance in fieldwork.

## Statement of inclusion

Our study brings together authors from different countries, including scientists based in the country where the study was carried out. All authors were engaged early on with the research and study design to ensure that the diverse sets of perspectives they represent was considered from the onset.

## Data availability statement

The data used in this study is available on request from the corresponding author.

## Supplementary materials

**Table S1.** Groups of Tree-related Microhabitats (TreMs) considered in the study with corresponding TreM types descriptions and catalogue codes (after Kraus et al. 2016).

| Group | Description / TreM types included | Catalogue codes |
| --- | --- | --- |
| Woodpecker cavities | Breeding cavities excavated by woodpeckers, $\varnothing = 4$ cm, 5-6 cm or $\varnothing > 10$ cm or at least three cavity openings in tree trunk within two meters | CV11, CV12, CV13, CV15 |
| Trunk and mould cavities with ground contact | Cavities with mould and ground contact, $\varnothing > 10$ cm or $\varnothing > 30$ cm | CV21, CV22 |
| Trunk and mould cavities without ground contact | Cavities with mould but without ground contact, $\varnothing > 10$ cm or $\varnothing > 30$ cm | CV23, CV24 |
| Large semi-open trunk and mould cavities | Semi-open or large trunk cavities with open top, $\varnothing > 30$ cm | CV25, CV26 |
| Branch holes | Rot-holes from branch breakage ($\varnothing > 5$ cm, $\varnothing > 10$ cm), hollow horizontal branches from breakage ($\varnothing > 10$ cm) | CV31, CV32, CV33 |
| Insect galleries and bore holes | Galleries with small ($< 2$ cm) or large ($> 2$ cm) bore holes | CV51, CV52 |
| Cracks and crevices | Injuries exposing cambium and sapwood, length $\geq 30$ cm or $\geq 100$ cm | IN31, IN32 |
| Bark pockets | Spaces between bark and sapwood, open at the bottom or at the top | BA11, BA12 |
| Cankers and burrs | Cankers without or with exposed necrotic tissue, $\varnothing > 20$ cm | GR31, GR32 |
| Canopy deadwood | Dead branches ($\varnothing 10\text{--}20$ cm or $> 20$ cm, $\geq 50$ cm length, sun-exposed or shaded) and dead tree tops ($\varnothing \geq 10$ cm) | DA11, DA12, DA13, DA14, DA15 |
| Fruiting bodies of fungi | Fruiting bodies of annual or perennial polypores, gill fungi, or large ascomycetes ($> 5$ cm) | EP11, EP12, EP13, EP14 |

